# microRNAs slow translating ribosomes to prevent protein misfolding

**DOI:** 10.1101/2020.12.08.417139

**Authors:** Hiroaki Sako, Takayuki Akimoto, Katsuhiko Suzuki, Takashi Ushida, Tadashi Yamamoto

## Abstract

An evolutionarily conserved mechanism, use of non-optimal codons, slows ribosomes during translation to allow proper folding of nascent polypeptides. However, until now, it was unknown whether any eukaryote-specific mechanisms exist for this purpose. Here, we propose that miRNAs slow translating ribosomes to prevent protein misfolding, with little negative effect on protein abundance. To prove this, we bioinformatically analyze ribosome profiling and miRNA binding sites and biochemically confirm that miRNA deficiency causes severe misfolding, which is rescued by slowing translating ribosomes. We demonstrate that non-cleaving shRNAs, targeting regions where elongation rates become faster in miRNA-deficient cells, improve protein folding with minimal effects on protein abundance. These results reveal broader functionality of miRNAs and a previously unknown mechanism to prevent protein misfolding.

**One Sentence Summary:** Eukaryote use of miRNAs prevents protein misfolding in a target-specific manner.

## Main Text

Proteins are sometimes misfolded during translation (*1*)(*2*). Even wild type *CFTR* (Cystic Fibrosis Transmembrane conductance Regulator gene), a mutation of which causes severe CFTR misfolding and cystic fibrosis, shows 80% misfolding rates during translation (*3*). To avoid such misfolding, there are evolutionarily conserved chaperon systems both in *E. coli* and humans. One of the chaperones, HSP70, is recruited to nascent polypeptides during translation to mediate correct folding (*4*)(*5*). Non-optimal codons are another evolutionarily conserved mechanism to effect proper folding (*6*)(*7*)(*8*). In both prokaryotes and eukaryotes, non-optimal codons, which recruit tRNAs slowly, reduce elongation rates, thereby allowing nascent proteins more time to associate with HSP70, to ensure proper folding (*4*)(*9*). Although these evolutionarily conserved mechanisms promote proper folding, we wondered if they are sufficient for the human proteome, which is much more complex than that of *E. coli*. Indeed, in degradation of proteins, eukaryotes have developed autophagy, which is not known among prokaryotes. As in degradation, it seemed likely that eukaryotes might have evolved additional mechanisms not seen in prokaryotes, so as to slow translating ribosomes where more precise folding is needed.

Most eukaryotes have microRNA (miRNA), but prokaryotes do not. miRNAs are believed to bind mainly to the 3’UTRs of mRNAs to trigger mRNA degradation or to inhibit translation. However, it is also true that most miRNAs target coding sequence (CDS) of mRNAs with highly diverse binding patterns and affinities, revealed by a series of studies in which the Ago-miRNA complex and its target sequence were ligated and co-purified, identifying which miRNAs bind to what parts of target sequences (*10*)(*11*)(*12*)(*13*). In view of these studies, we speculated that miRNA could be a eukaryote-specific mechanism to prevent protein misfolding during translation. Here, we hypothesize that miRNAs transiently slow active ribosomes to enhance nascent protein folding.

### Weakly binding miRNAs briefly pause translating ribosomes with little effect on translation

From a bioinformatic perspective, we first sought to determine whether miRNAs can slow actively translating ribosomes without strong translation inhibition. To this end, we conducted ribosome profiling and mRNA-seq, as well as prediction of miRNA binding sites on CDSs (Fig. 1A, S1A). There is often noise in meta-gene analysis of ribosomal dwell time (Fig. S1B, C). We assumed such noise comes from expression diversity of highly and less highly translated genes. Many factors in mRNA (e.g., secondary structure, RNA binding proteins, codon optimality, and miRNA) affect ribosomal dwell time at each codon. Small portions of highly translated genes, biased in such factors, mask the majority of less highly expressed genes, which makes noiseless analysis difficult. To circumvent this obstacle, we equalized contributions from highly and rarely translated genes and confirmed the robustness of our noise reduction method (Fig. S1B, C). Using this method, we then calculated the relative distance of ribosome footprints from miRNAs and their densities, in order to analyze the effect of miRNAs on translating ribosomes (Fig. 1B, S1D). There were 3 features common to both mouse skeletal muscle and HEK293 analyses. 1) a ribosomal footprint peak exists at 0 nt of the relative distance, regardless of miRNA binding strength, suggesting that weakly binding miRNAs can also slow translating ribosomes. 2) There is a dip in ribosome density at +20 to +25 nt only for moderately and strongly binding miRNAs. This indicates that there are fewer translating ribosomes downstream of moderately and strongly binding miRNAs, suggesting that such miRNAs slow ribosomes more effectively. 3) Overall ribosome densities are higher with weakly binding miRNAs compared to those that bind more strongly, indicating that translation inhibition is less likely with weakly binding miRNAs. Taken together, this bioinformatic analysis supports the idea that weakly binding miRNAs slow translating ribosomes with little effect on translation.

**Fig. 1.**
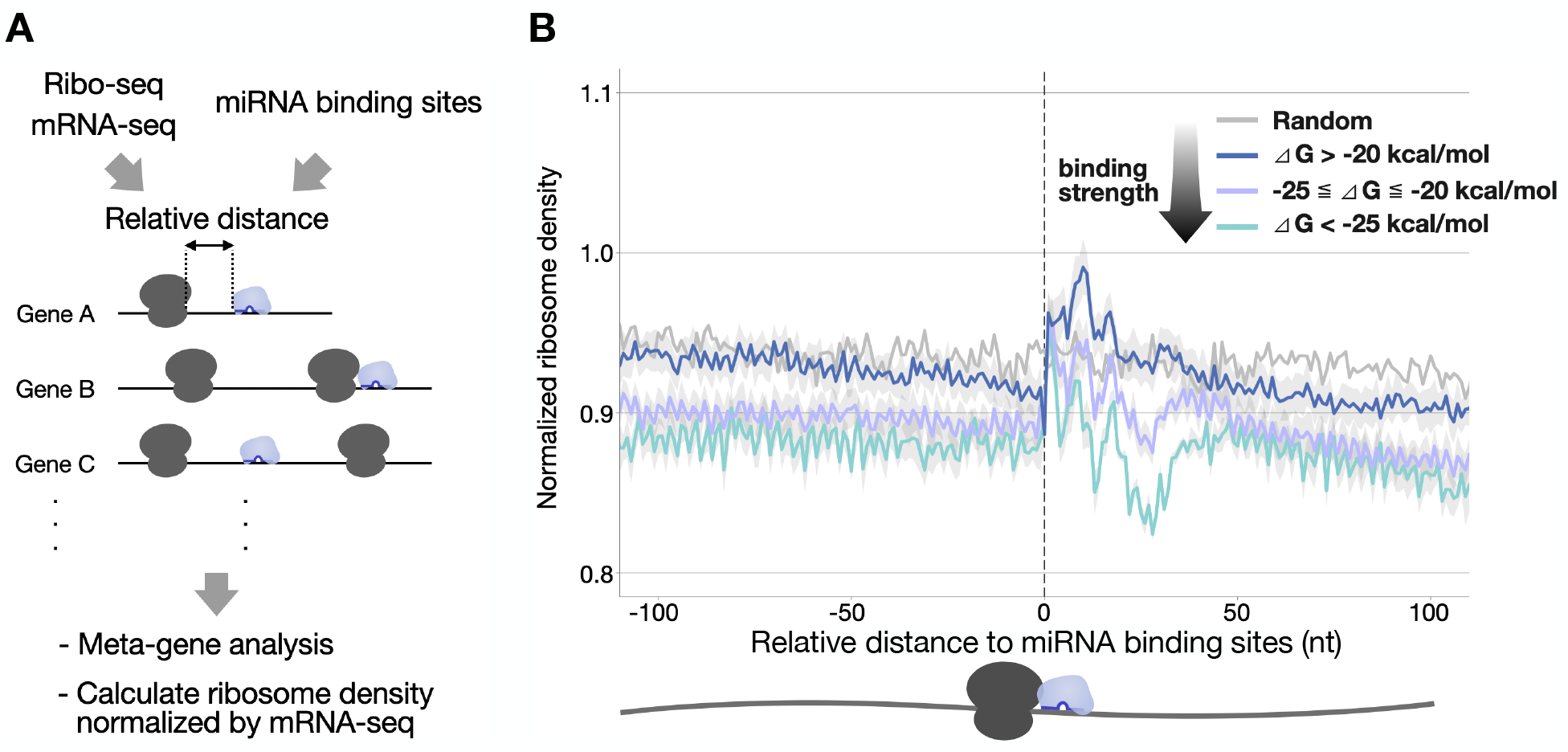
Weakly binding miRNAs on CDSs slow translating ribosomes with little effect on translation efficiency. (A) Schematic image of the meta-gene analysis to show ribosome densities relative to predicted miRNA binding sites on CDSs. (B) Normalized ribosome densities are shown. 3’ ends of ribosome footprints relative to the 5’ ends of miRNA binding sites were calculated to show accumulated ribosome densities normalized by mRNA-seq. Lines of different colors indicate different miRNA binding strengths. N=3. Mean ± SE.

### Fewer miRNAs, more NBD1 misfolding

To experimentally verify the model, we designed misfolding-prone reporters (Fig 2A) composed of EGFP-tagged NBD1 (Nucleotide-Binding Domain 1) of CFTR and a P2A sequence to separately express mCherry. This NBD1 domain is highly vulnerable to misfolding during translation (*3*). For miRNA-depleted cells, miRNA processing enzymes, DROSHA or AGO1 and 2, were targeted with the CRISPR/Cas9 system, denominated sgDROSHA and sgAGO1/2, respectively (Fig. S2A-C). Based on our model, we expected that reduced miRNA would enhance misfolding and aggregation of NBD1. Indeed, proteasome inhibition formed more NBD1 perinuclear puncta in sgDROSHA and sgAGO1/2 compared to the NTC control (Fig 2B). This was confirmed with NBD1 immunoblots. More insoluble NBD1 was detected in sgDROSHA and sgAGO1/2 insoluble fractions (Fig. 2C, S2E), but not in total extracts (Fig. 2D, S2F). Although these data indicate that fewer miRNAs lead to more misfolding of NBD1, we were still not sure whether the observed NBD1 misfolding was attributable to misfolding of newly synthesized NBD1 or to misfolding of pre-existing mature NBD1.

**Fig. 2.**
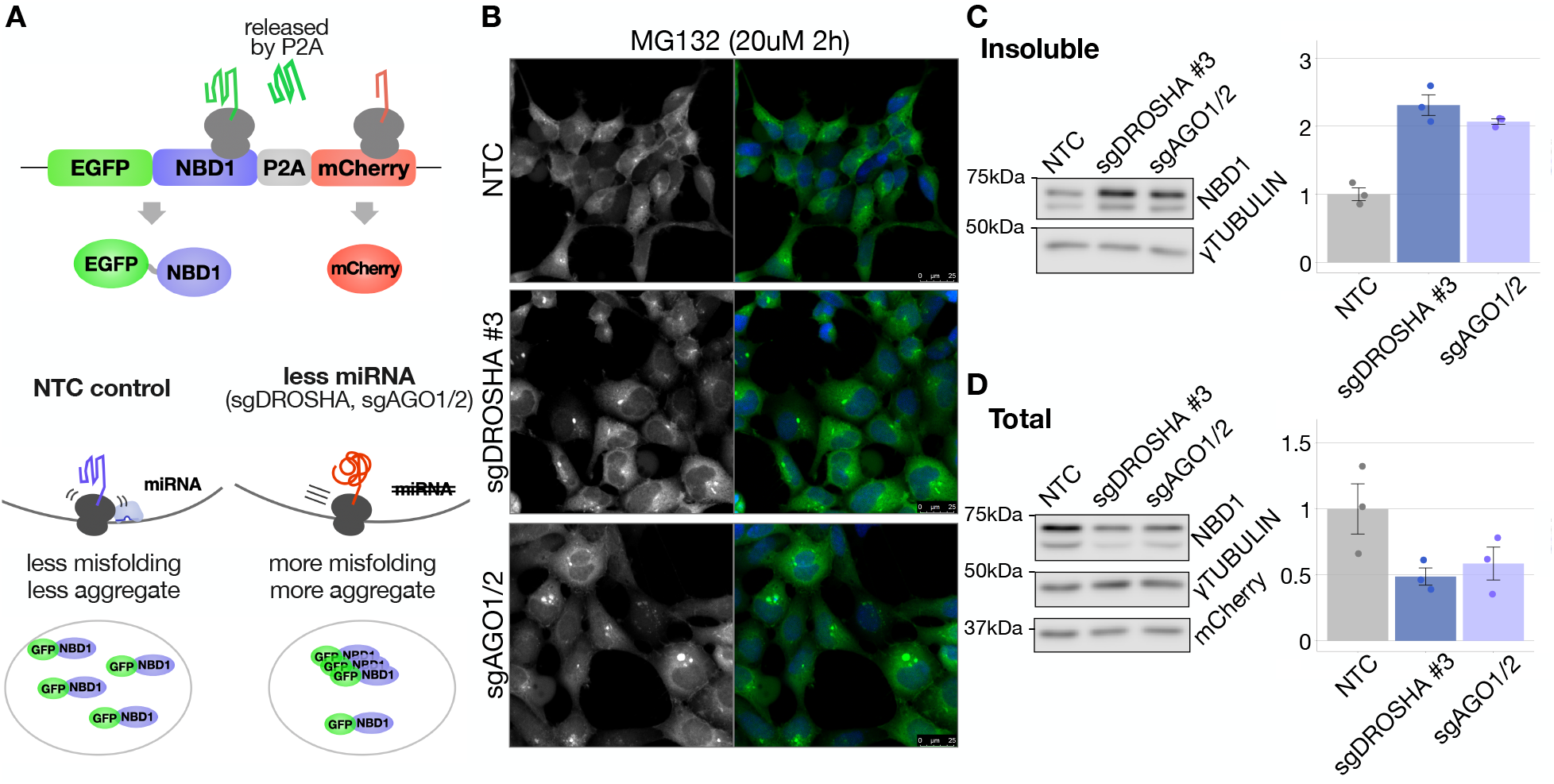
Fewer miRNAs, more NBD1 misfolding. (A) Schematic image of the misfolding-prone NBD1 reporter and a proposed working model. (B) Confocal images showing aggregations of misfolded EGFP-tagged NBD1 reporter in NTC control, DROSHA knockout, or AGO1 and 2 double-knockout cells. Nuclei were counterstained blue with Hoechst. (C) Immunoblots and quantification of insoluble protein fractions of NBD1 after 120 min of MG132 (20 μM) treatment, normalized against γTUBULIN for the plot. The NBD1 blot shows two bands, each cleaved by 1^st^ P2A or 2^nd^ P2A sequences present upstream of the EGFP-NBD1 sequence. (D) Total extract of (C). NBD1 was normalized against γTUBULIN for the plot. N=3. Mean ± SE.

### Translation slowdown rescues NBD1 misfolding without inhibiting its translation

Our model predicts that miRNA slows translating ribosomes to prevent misfolding of nascent peptides. Thus, slowing of translation should prevent NBD1 misfolding only if the misfolding originates with nascent NBD1, not from mature protein. To prove this, we slowed elongating ribosomes with very low concentrations of a reversible translation inhibitor, cycloheximide (CHX), to see whether CHX could rescue NBD1 misfolding without affecting translation efficiency. As expected, CHX (0.1 μg/mL) rescued NBD1 misfolding caused by sgDROSHA or sgAGO1/2 (Fig. 3A, B) without affecting total protein or NBD1 reporter synthesis rates (Fig. 3C, S3C, D). These observations confirm that reducing the speed of translation rescues nascent NBD1 misfolding caused by reduced levels of miRNAs.

**Fig. 3.**
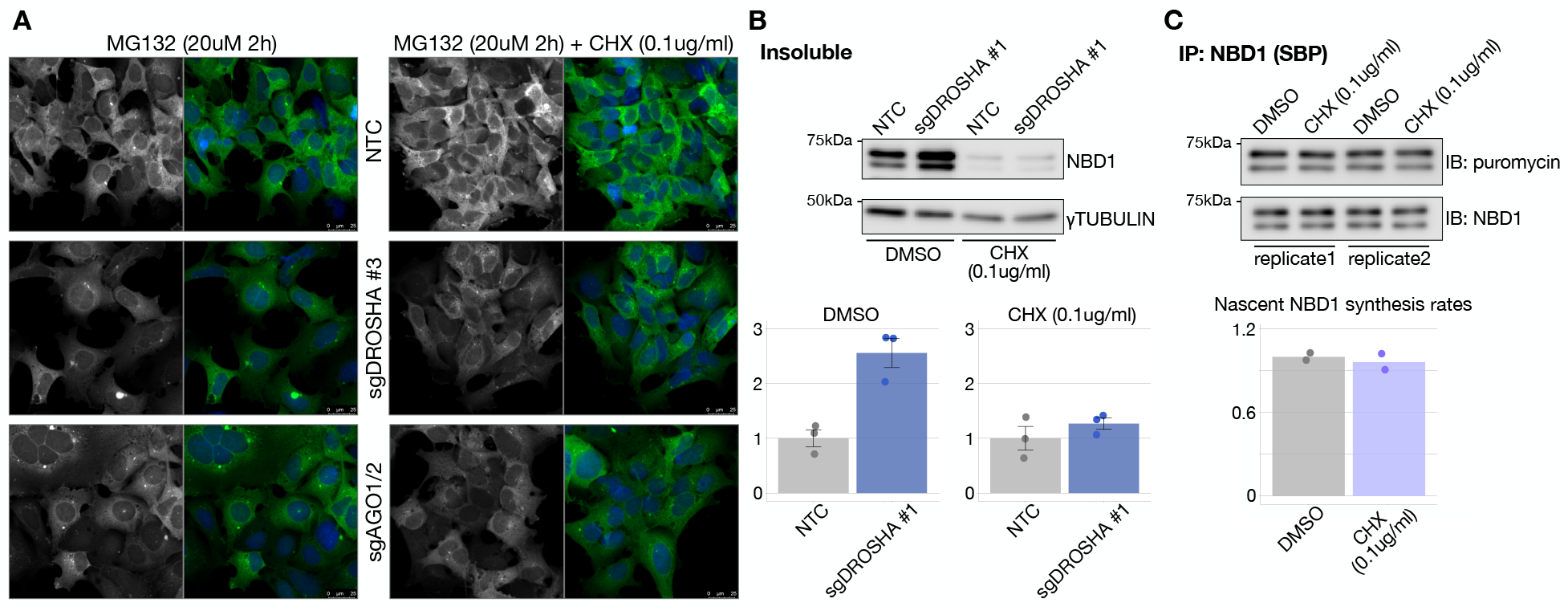
Slowing of translation rescues NBD1 misfolding without inhibiting its translation. (A) Confocal images showing aggregations of misfolded EGFP-tagged NBD1 reporter with or without a very low concentration of cycloheximide (CHX). Nuclei were counterstained blue with Hoechst. (B) Immunoblots and quantification of insoluble protein fractions with or without CHX. The NBD1 reporter was normalized against γTUBULIN for the plot. N=3. Mean ± SE. (C) NBD1 reporter-specific SUnSET assay. Nascent NBD1 reporter detected with anti-puromycin was normalized against immunoprecipitated NBD1 reporter. N=2.

### Fewer miRNAs, less co-translational recruitment of HSC70

Although reducing translation speed rescues NBD1 misfolding induced by miRNA deficiency, we were still not sure whether reduced miRNA levels actually accelerate elongating ribosomes. If so, sgDROSHA should decrease co-translational recruitment of HSC70, a constitutive form of HSP70 associated with CFTR maturation (*14*)(*15*). To test this hypothesis, we carried out RIP-qPCR (RNA Immunoprecipitation qPCR), targeting HSC70 for immunoprecipitation and an NBD1 reporter for qPCR. As hypothesized, sgDROSHA reduced HSC70 recruitment to nascent NBD1 compared to NTC controls (Fig 4A, B, S4B). Importantly, reduced HSC70 recruitment was not caused by a decrease in total HSC70 abundance or in HSC70-NBD1 interaction efficiency, because total HSC70 was unchanged (Fig. 4A) and the interaction efficiency of HSC70 and mature NBD1 was also unchanged (Fig. S4C). Moreover, the result was not caused by reduced NBD1 translation efficiency (mRNA-seq showed *NBD1* expression ratio of NTC : sgDROSHA = 1.00 : 1.00 and ribosome profiling showed NTC : sgDROSHA = 1.00 : 0.93). These results suggest that reduced levels of miRNAs cause inefficient co-translational recruitment of HSC70.

**Fig. 4.**
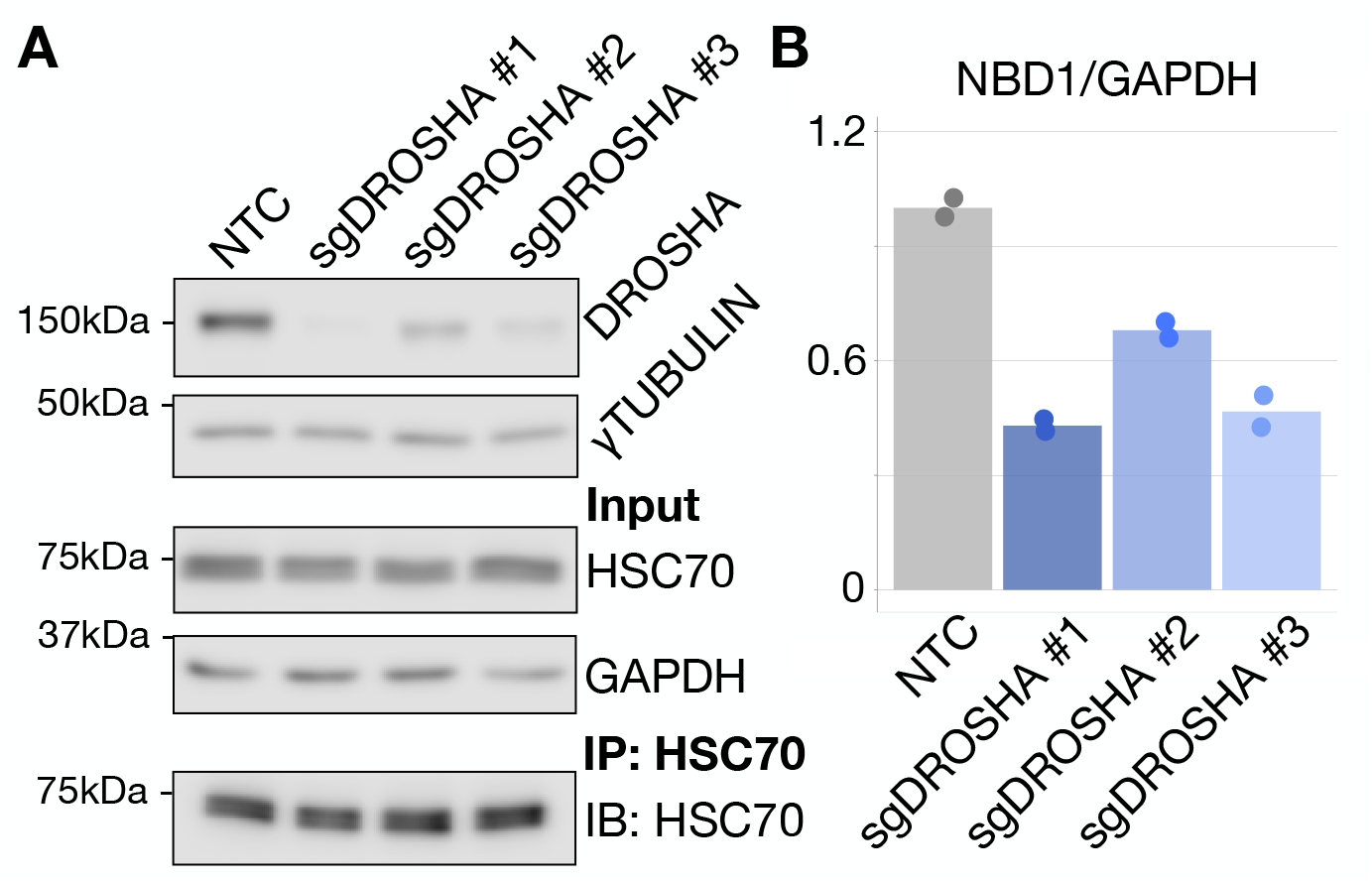
Fewer miRNAs, less co-translational recruitment of HSC70. (A) Immunoblots of input and IP samples. (B) RIP-qPCR. Quantification of *NBD1 reporter* mRNA levels, normalized against *GAPDH*. IP: HSC70. N=2.

### Buffering of NBD1-specific miRNAs causes more NBD1 misfolding

However, global depletion of miRNAs has many primary and secondary effects. It is possible that observed results were actually caused by loss of conventional miRNA functions, rather than by loss of miRNA-induced ribosomal pausing. To minimize conventional effects of miRNAs, we attempted to buffer miRNAs targeting NBD1 sequences (i.e., reducing miRNA-ribosome collision rates in a NBD1 specific manner). To this end, we prepared another NBD1 reporter, stop-NBD1, which expresses more NBD1 transcripts, but translates at lower efficiency (Fig. S6A, B). These extra NBD1 transcripts can buffer miRNAs targeting NBD1. We assumed that lower translation efficiency leads to lower miRNA-ribosome collision rates, which makes NBD1 more prone to misfolding. In fact, stop-NBD1 resulted in a higher misfolding ratio compared to the higher translation efficiency of the NBD1 reporter (Fig. S6C), further supporting our hypothesis. However, so far, we have only shown strong “associations” between miRNA deficiency and misfolding. At this point, there are still some deficiencies in our proof. 1) Evidence of accelerated elongation in miRNA-depleted cells is still lacking. 2) Causality between miRNA and misfolding prevention is still missing.

### shRNA reduces nascent protein misfolding with a minimal effect on translation

To examine the effect of miRNA reduction on translating ribosomes and to demonstrate causality between miRNA and prevention of misfolding, we performed ribosome profiling using sgDROSHA cells and designed shRNA targeting the region where sgDROSHA accelerates elongation rates. Because cellular mechanisms to process shRNA largely overlap with those for miRNA, we consider shRNA as the best choice to mimic miRNA. In ribosome profiling, ribosome density is expected to be lower if elongation rates are accelerated in sgDROSHA cells. We found a 60-nt region in the NBD1 domain that reproducibly showed lower ribosome densities in sgDROSHA compared to NTC controls (Fig. 5A, S7A). Thus, this 60-nt sequence was the primary target of shRNA. If accelerated elongation in the region is responsible for NBD1 misfolding in miRNA-deficient cells, non-cleaving shRNAs targeting the region should improve NBD1 folding even in NTC controls. Indeed, tandemly expressed shRNAs (especially 1-1M/4-0M and 1-2M/4-2M) reduced NBD1 misfolding with minimal effect on total NBD1 abundance (Fig 5B, C, S7D). Note that mCherry in total extracts decreased (Fig. 5C), implying translational inhibition. However, the decrease in total NBD1 was not as much as in mCherry (Fig. 5C, 1-1M/4-0M). This suggests that even though NBD1 and mCherry translation is inhibited to some extent by shRNA, shRNA-mediated pausing/folding also stabilizes nascent NBD1, recovering total NBD1 abundance. However, this is not the case for mCherry. Collectively, with minimal effects on total NBD1 abundance, NBD1 folding was improved by shRNA targeting sequences where translation elongation accelerated in sgDROSHA, supporting a model that miRNAs slow translating ribosomes to prevent misfolding, with minimal effect on translation.

**Fig. 5.**
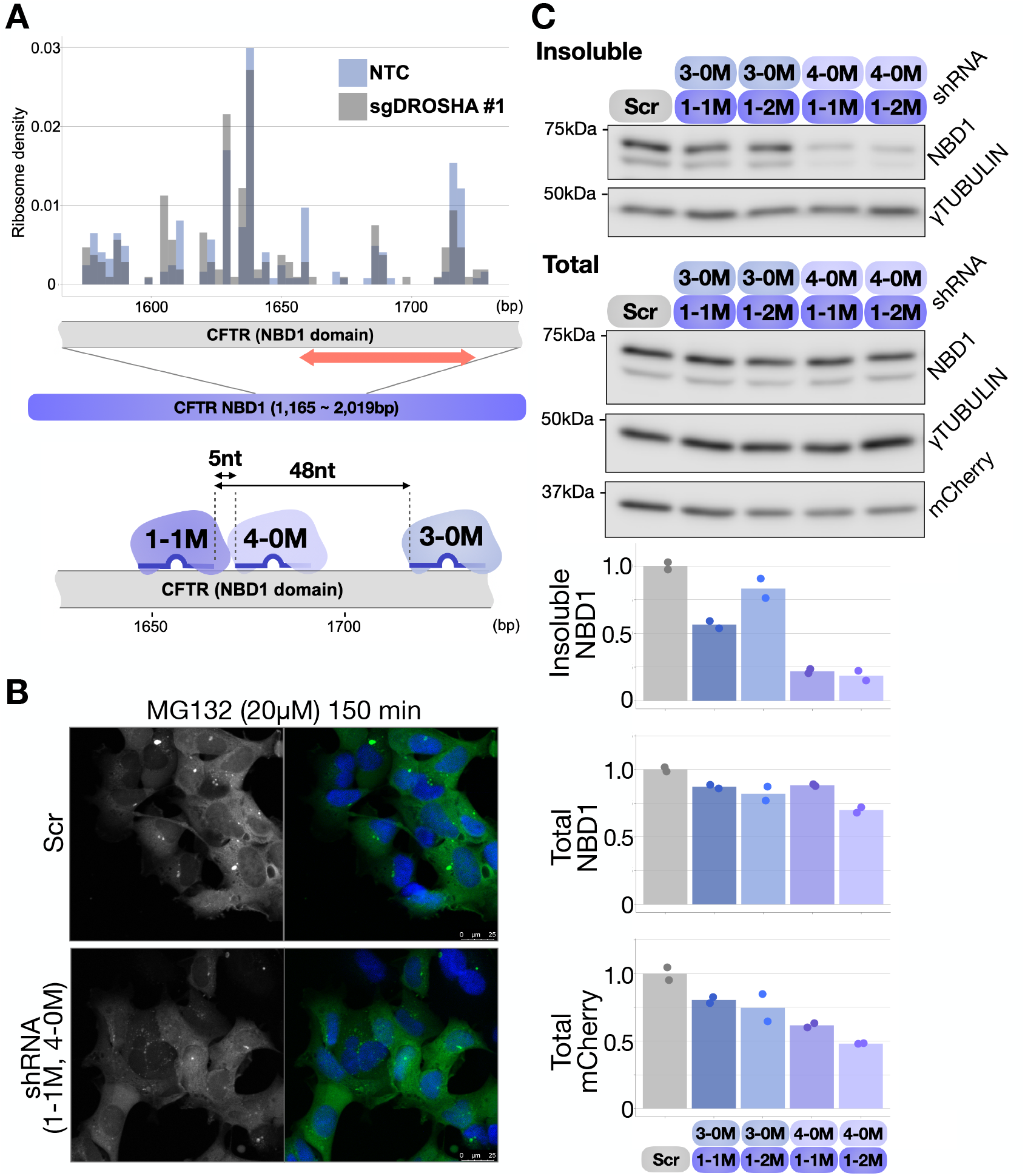
shRNAs prevent NBD1 misfolding with minimal effect on total NBD1. (A) Ribosome densities of the NBD1 domain in NTC and sgDROSHA are shown. Raw ribosomal footprint counts were normalized against total footprint numbers aligned to the NBD1 domain sequence. Bin = 3 nt, showing single-codon resolution. The target (∼ 60 nt) of shRNA is indicated by a red arrow. At the bottom is a schematic image of shRNAs targeting the 60-nt region. shRNA “4-0M” indicates shRNA #4 which has no mismatch at the 3’ end of the shRNA. All shRNA used here is non-cleaving. (B) Confocal images showing aggregations of misfolded EGFP-tagged NBD1 reporter in the Scr control and tandemly expressed shRNAs (1-1M/4-0M). Nuclei were counterstained blue with Hoechst. (C) Immunoblots and quantification of insoluble fractions of NBD1 and total NBD1 and mCherry after 150 min of MG132 (20 μM) treatment, normalized against γTUBULIN for the plot. N=2.

**Fig. 6.**
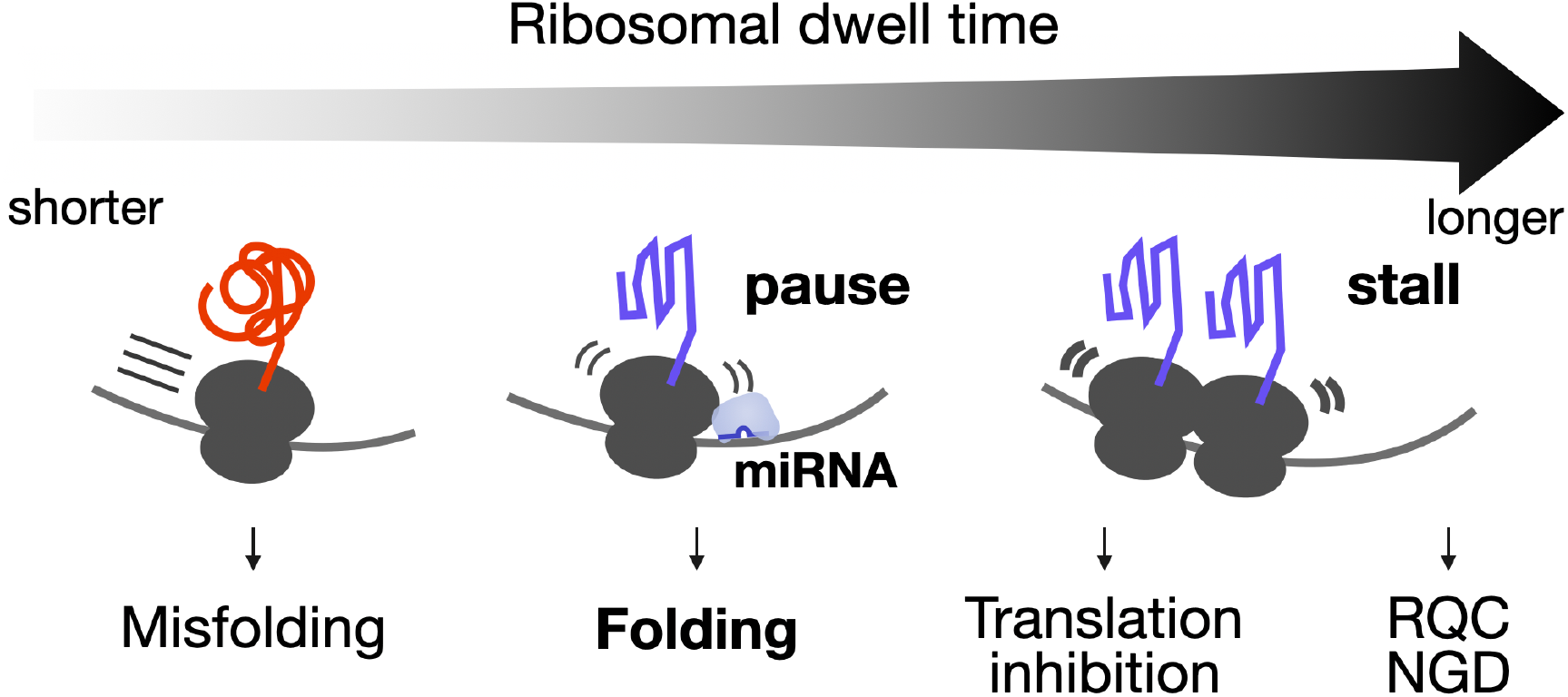
Schematic image of the working model. Faster translation (shorter ribosomal dwell time) leads to nascent peptide misfolding. Ribosomal stall (longer dwell time) induces translation inhibition or RQC/NGD. The dwell time of miRNA-mediated pausing seems moderate.

## Discussion

We discovered that miRNAs slow translating ribosomes and enhance proper polypeptide folding during translation. Ribosomal dwell time seems important for miRNA-mediated pausing/folding. As far as the misfolding-prone domain is concerned, too short a dwell time or too rapid elongation tends to exacerbate misfolding (*16*)(*3*)(*2*). In contrast, if dwell time becomes too long, ribosomes collide and form disomes, triggering translation shutoff (*17*) or RQC (Ribosome-associated Quality Control) and NGD (No-Go Decay) pathways (*18*)(*19*)(*20*)(*21*). Thus, ribosomal dwell time must be neither too short nor too long during miRNA-mediated pausing. We estimate average pausing time without a collision up to ∼10 seconds given that there is one translating ribosome per ∼ 135 nt in sea urchins (*22*) and 189 nt in 293T cells (*23*) and that the average translating ribosome rate is ∼ 5.6 amino acid per second (*24*).

Although we suggested that recruitment of HSC70 is a key to miRNA-mediated folding, mechanistic details remain largely unknown. Since tandemly expressed shRNAs (Fig. 5), TNRC6A (GW182), tethering up to three AGOs (*25*), may play a role in modulating binding strength and/or stability of steric hindrance to slow ribosomes appropriately. It will also be intriguing to see whether miRNA-mediated pausing must avoid ribosomal collision detectors, such as ZAKα (*17*), ZNF598 (Hel2) (*21*), and EDF1 (*26*)(*27*) or how miRNA-mediated pausing interacts with a non-optimal codon monitor, CCR4-NOT complex (*28*).

We used a reporter system containing a CFTR NBD1 domain. miRNA-mediated NBD1 stabilization could be a previously unknown approach to stabilize CFTR proteins in cystic fibrosis patients. Finally, other proteins could be targeted by miRNA-mediated pausing/folding. Considering that non-optimal codons have a significant role in efficient recognition of signaling peptides (*29*)(*6*), miRNA-mediated pausing/folding may prevent mis-translocation of mitochondrial proteins, or it may enhance secreted protein production, such as insulin in diabetes.

## Acknowledgments

We thank Tsukasa Okiyoneda for discussion and advice. We thank Steven D. Aird for editing the manuscript (www.sda-technical-editor.org).

## Author contributions

HS designed and performed experiments and analyzed data. All authors discussed the results and wrote the manuscript.

## Data and materials availability

Sequencing data have been deposited in the National Center for Biotechnology Information’s Gene Expression Omnibus under accession number GSE160917.

## Supplementary Materials

Materials and Methods

Figures S1-S7

Tables S1

References (*30-43*)

## References

1. C. Kimchi-Sarfaty, J. M. Oh, I.-W. Kim, Z. E. Sauna, A. M. Calcagno, S. V. Ambudkar, M. M. Gottesman, A “Silent” Polymorphism in the MDR1 Gene Changes Substrate Specificity. Science. 315, 525–528 (2007).

2. M. Zhou, J. Guo, J. Cha, M. Chae, S. Chen, J. M. Barral, M. S. Sachs, Y. Liu, Non-optimal codon usage affects expression, structure and function of clock protein FRQ. Nature. 495, 111–115 (2013).

3. S. J. Kim, J. S. Yoon, H. Shishido, Z. Yang, L. A. Rooney, J. M. Barral, W. R. Skach, Translational tuning optimizes nascent protein folding in cells. Science. 348, 444–448 (2015).

4. K. C. Stein, A. Kriel, J. Frydman, Nascent Polypeptide Domain Topology and Elongation Rate Direct the Cotranslational Hierarchy of Hsp70 and TRiC/CCT. Mol. Cell. 75, 1117-1130.e5 (2019).

5. K. Döring, N. Ahmed, T. Riemer, H. G. Suresh, Y. Vainshtein, M. Habich, J. Riemer, M. P. Mayer, E. P. O’Brien, G. Kramer, B. Bukau, Profiling Ssb-Nascent Chain Interactions Reveals Principles of Hsp70-Assisted Folding. Cell. 170, 298-311.e20 (2017).

6. S. Pechmann, J. W. Chartron, J. Frydman, Local slowdown of translation by nonoptimal codons promotes nascent-chain recognition by SRP in vivo. Nat. Struct. Mol. Biol. 21, 1100–1105 (2014).

7. K. C. Stein, J. Frydman, The stop-and-go traffic regulating protein biogenesis: How translation kinetics controls proteostasis. J. Biol. Chem. 294, 2076–2084 (2019).

8. J. L. Chaney, A. Steele, R. Carmichael, A. Rodriguez, A. T. Specht, K. Ngo, J. Li, S. Emrich, P. L. Clark, Widespread position-specific conservation of synonymous rare codons within coding sequences. PLOS Comput. Biol. 13, e1005531 (2017).

9. P. Cortazzo, C. Cerveñansky, M. Mariń, C. Reiss, R. Ehrlich, A. Deana, Silent mutations affect in vivo protein folding in Escherichia coli. Biochem. Biophys. Res. Commun. 293, 537–541 (2002).

10. A. Helwak, G. Kudla, T. Dudnakova, D. Tollervey, Mapping the Human miRNA Interactome by CLASH Reveals Frequent Noncanonical Binding. Cell. 153, 654–665 (2013).

11. S. Grosswendt, A. Filipchyk, M. Manzano, F. Klironomos, M. Schilling, M. Herzog, E. Gottwein, N. Rajewsky, Unambiguous Identification of miRNA:Target Site Interactions by Different Types of Ligation Reactions. Mol. Cell. 54, 1042–1054 (2014).

12. M. J. Moore, T. K. H. Scheel, J. M. Luna, C. Y. Park, J. J. Fak, E. Nishiuchi, C. M. Rice, R. B. Darnell, miRNA–target chimeras reveal miRNA 3′-end pairing as a major determinant of Argonaute target specificity. Nat. Commun. 6, 8864 (2015).

13. J. P. Broughton, M. T. Lovci, J. L. Huang, G. W. Yeo, A. E. Pasquinelli, Pairing beyond the Seed Supports MicroRNA Targeting Specificity. Mol. Cell. 64, 320–333 (2016).

14. G. C. Meacham, The Hdj-2/Hsc70 chaperone pair facilitates early steps in CFTR biogenesis. EMBO J. 18, 1492–1505 (1999).

15. I. Baaklini, C. de C. Gonçalves, G. L. Lukacs, J. C. Young, Selective Binding of HSC70 and its Co-Chaperones to Structural Hotspots on CFTR. Sci. Rep. 10, 4176 (2020).

16. F. Buhr, S. Jha, M. Thommen, J. Mittelstaet, F. Kutz, H. Schwalbe, M. V. Rodnina, A. Komar, Synonymous Codons Direct Cotranslational Folding toward Different Protein Conformations. Mol. Cell. 61, 341–351 (2016).

17. C. C.-C. Wu, A. Peterson, B. Zinshteyn, S. Regot, R. Green, Ribosome Collisions Trigger General Stress Responses to Regulate Cell Fate. Cell. 182, 404-416.e14 (2020).

18. M. H. Bengtson, C. A. P. Joazeiro, Role of a ribosome-associated E3 ubiquitin ligase in protein quality control. Nature. 467, 470–473 (2010).

19. T. Tsuboi, K. Kuroha, K. Kudo, S. Makino, E. Inoue, I. Kashima, T. Inada, Dom34:Hbs1 Plays a General Role in Quality-Control Systems by Dissociation of a Stalled Ribosome at the 3′ End of Aberrant mRNA. Mol. Cell. 46, 518–529 (2012).

20. O. Brandman, J. Stewart-Ornstein, D. Wong, A. Larson, C. C. Williams, G.-W. Li, S. Zhou, D. King, P. S. Shen, J. Weibezahn, J. G. Dunn, S. Rouskin, T. Inada, A. Frost, J. S. Weissman, A Ribosome-Bound Quality Control Complex Triggers Degradation of Nascent Peptides and Signals Translation Stress. Cell. 151, 1042–1054 (2012).

21. C. L. Simms, L. L. Yan, H. S. Zaher, Ribosome Collision Is Critical for Quality Control during No-Go Decay. Mol. Cell. 68, 361-373.e5 (2017).

22. K. A. Martin, O. L. Miller, Polysome structure in sea urchin eggs and embryos: An electron microscopic analysis. Dev. Biol. 98, 338–348 (1983).

23. D. G. Hendrickson, D. J. Hogan, H. L. McCullough, J. W. Myers, D. Herschlag, J. E. Ferrell, P. O. Brown, Concordant Regulation of Translation and mRNA Abundance for Hundreds of Targets of a Human microRNA. PLoS Biol. 7, e1000238 (2009).

24. N. T. Ingolia, L. F. Lareau, J. S. Weissman, Ribosome Profiling of Mouse Embryonic Stem Cells Reveals the Complexity and Dynamics of Mammalian Proteomes. Cell. 147, 789–802 (2011).

25. E. Elkayam, C. R. Faehnle, M. Morales, J. Sun, H. Li, L. Joshua-Tor, Multivalent Recruitment of Human Argonaute by GW182. Mol. Cell. 67, 646-658.e3 (2017).

26. N. K. Sinha, A. Ordureau, K. Best, J. A. Saba, B. Zinshteyn, E. Sundaramoorthy, A. Fulzele, D. M. Garshott, T. Denk, M. Thoms, J. A. Paulo, J. W. Harper, E. J. Bennett, R. Beckmann, R. Green, EDF1 coordinates cellular responses to ribosome collisions. eLife. 9, e58828 (2020).

27. S. Juszkiewicz, G. Slodkowicz, Z. Lin, P. Freire-Pritchett, S.-Y. Peak-Chew, R. S. Hegde, Ribosome collisions trigger cis-acting feedback inhibition of translation initiation. eLife. 9, e60038 (2020).

28. R. Buschauer, Y. Matsuo, T. Sugiyama, Y.-H. Chen, N. Alhusaini, T. Sweet, K. Ikeuchi, J. Cheng, Y. Matsuki, R. Nobuta, A. Gilmozzi, O. Berninghausen, P. Tesina, T. Becker, J. Coller, T. Inada, R. Beckmann, The Ccr4-Not complex monitors the translating ribosome for codon optimality. Science. 368, eaay6912 (2020).

29. J. W. Chartron, K. C. L. Hunt, J. Frydman, Cotranslational signal-independent SRP preloading during membrane targeting. Nature. 536, 224–228 (2016).

