## Supplementally materials for "microRNAs slow translating ribosomes to prevent protein misfolding"

**This PDF file includes:**

Materials and Methods
Figs. S1 to S7
Tables S1

**Materials and Methods**

Animals

Seven-week-old male C57BL/6J mice, purchased from Kiwa Laboratory Animals (Wakayama, Japan), were first acclimated to the lab environment for a week (temperature controlled at 23 °C with 12 hr light/ dark cycle and fed with normal chow).

Plasmid construction

For misfolding-prone NBD1 reporters targeting the human CFTR (NM_000492.4), the NBD1 domain (T389 to G673) was PCR-amplified (sequence was used to confirm that Val470 was mutated to Met, identical to AH006034.2) with a linker (GGGGS) at the 5′ end and (linker)x2 (GGGGSGGGGS) at the 3′ end. NBD1 was then inserted to pMX-RGb, a gift from Yoshio Kato (*30*). SBP tag and P2A sequence was added between the downstream of the NBD1 and mCherry-Flag sequences and Flag-EGFP was inserted upstream of the NBD1 domain. The reporter plasmid, pMX-Flag-EGFP-linker-NBD1-(linker)x2-SBP-P2A-mCherry-Flag, named “gag-pol-NBD1 reporter”, yielding gag-pol-fused NBD1 and gag-pol free NBD1. This is because there is a relatively weak Kozac sequence at the gag-pol translation start site. Ribosomes scanning through the gag-pol translation start site initiate translation at Flag. To separately produce gag-pol and Flag-NBD1-EGFP-SBP, double P2A sequences were inserted between the pol and Flag sequences, called an “NBD1 reporter.” The NBD1 reporter expresses two different lengths of Flag-EGFP-NBD1-SBP cleaved by the 1$^{st}$ P2A or 2$^{nd}$ P2A sequences as seen in the immunoblots. A further modified reporter, in which double P2A between pol and Flag was replaced with 4 stop codons, was prepared and named “stop-NBD1 reporter” to express Flag-EGFP-NBD1-SBP at lower translation efficiency compared to the NBD1 reporter.

For the CRISPR/Cas9 system, guide sequences (Invitrogen) were designed using Benchling (https://benchling.com/) as listed in Supplementary Table S1. Guide oligos were 5′ phosphorylated, annealed, and ligated to BsmBI (NEB)-digested lenti-CRISPR V2 (*31*) (a gift from Feng Zhang, Addgene #52961).

shRNA-expressing plasmids were prepared as follows. For non-cleaving shRNA, shRNA sequences were designed to have 2 mismatches at the 10$^{th}$ and 11$^{th}$ bases to avoid AGO-shRNA-induced cleavage of the target mRNA. To tandemly express shRNA, shRNA top and bottom sequences were 5′ phosphorylated, annealed, and ligated to BglII-XhoI-digested pSuperior (oligoengine) in which a SalI digestion site upstream of the H1 promoter was inserted in advance. Then, shRNA together with the H1 promoter were SalI-XhoI-digested to ligate to SalI-digested pLKO.1-puro (*32*) (a gift from Bob Weinberg, Addgene #8453) in which the U6 promoter was replaced with a SalI digestion site in advance. Because SalI and XhoI are palindromic, SalI-XhoI-digested H1-shRNA can be ligated to SalI-digested pLKO.1-puro. Similarly for a 2nd shRNA insertion, SalI-digested pLKO.1-puro-shRNA (1st) can be ligated to a SalI-XhoI-digested 2nd shRNA, which can tandemly express 1st and 2nd shRNAs.

<u>Cell lines and culture conditions</u>
All cell lines were cultured in DMEM with high glucose (Fujifilm, Japan) with 10% FBS (Nichirei, Japan) and 1% penicillin/streptomycin.

gag-pol-NBD1, stop-NBD1, or NBD1 reporters and pCMV-VSVg (*32*) (a gift from Bob Weinberg, Addgene #8454) were co-transfected to PlatA (Cell Biolabs) using Lipofectamine 3000 (Invitrogen). Two days after transfection, supernatant was filtered through a 0.45 µm pore filter and centrifuged at 20,000 g at 4 °C for 5hr. Pellets were resuspended with DMEM culture medium in which 293T cells (American Type Culture Collection) were infected for 2 days. Since the translation efficiency of the stop-NBD1 reporter is lower than the others, stop-NBD1 was infected at 3 to 4 fold higher titer.

The CRISPR/Cas9 system was used to prepare AGO1 and AGO2 double-knockout or DROSHA-knockout cells. Lenti-CRISPR V2 (NTC, sgDROSHA, or sgAGO1 and sgAGO2), pCMV-VSVg, and psPAX2 (a gift from Didier Trono, addgene #12260) were co-transfected to PlatA. Two days after transfection, supernatant (10 mL) was filtered through a 0.45 µm pore filter and diluted with fresh 10 mL DMEM with polybrene (6 µg/mL) in which gag-pol-NBD1 or NBD1 reporter clonal cell lines were infected for 2 days, followed by 2 µg/mL puromycin selection.

For human AGO2 (5xA), five phosphorylation sites in the PIWI domain (S824 to S834) were converted to alanine, which enhances the targeting of miRNA to CDS compared to WT (*33*), sgAGO1/2 NBD1 reporter cells were infected with either an empty vector or human AGO2 (5xA) (gifts from Joshua Mendell, Addgene #91980 and Addgene #91979, respectively) (*33*) as described above. Then, a clonal 293T cell line stably expressing NBD1 reporter was infected with the resulting shRNA, as described above.

<u>Ribosome profiling and mRNA-seq</u>
Ribosome profiling protocols were modified from the originals (*34*)(*24*) and mouse skeletal muscle (*35*) and a TruSeq Ribo Profile (Mammalian) Kit (Illumina) were used according to the manufacturer's instructions with some modifications.

For mouse skeletal muscle samples, five µL of 50 mg/mL cycloheximide (Sigma) in PBS were injected into the right gastrocnemius muscles of mice (aged 9 to 10 weeks) and the right gastrocnemius was immediately harvested after mice were sacrificed. Harvested samples were washed in 100 µg/mL cycloheximide in PBS and flash-frozen in liquid nitrogen. Cycloheximide-treated samples were further processed with mRNA-seq and ribosome profiling as described below. Muscle samples were homogenized in 1.2 mL of ice-cold lysis buffer (20 mM Tris HCl pH 7.4, 150 mM NaCl, 5 mM $MgCl_2$, 1 mM DTT, 100 µg/mL cycloheximide, TURBO DNase I 25 U/mL, and 1% Triton X-100), followed by centrifugation at 15,000 g for 10 min at 4 °C. Supernatant from the lysate was separated into two 600 µL aliquots, one for mRNA-seq and the other for ribosome profiling.

For 293T cells, cells were not pre-treated with cycloheximide, but were washed in 100 µg/mL cycloheximide in PBS and lysed in 600 µL ice-cold lysis buffer containing 100 µg/mL cycloheximide.

For ribosome profiling, lysate was digested with RNase-If (10 unit/μg RNA) (NEB) for 45 min at room temperature, followed by addition of 15 μL SuperaseIn (Invitrogen). Three hundred μL lysate were used for sucrose cushion (900 μL sucrose cushion buffer, 100,000 rpm at 4 °C for 2 hr with TLA100.3 rotar [Beckman Coulter], Optima TLX [Beckman Coulter]), followed by PAGE purification (26 ~ 30 nt) and end-repair. For mRNA-seq, polyA RNA was extracted from total RNA using a Dynabeads mRNA DIRECT Micro Kit (Life technologies) and fragmented at 95 °C for 25 min, followed by end-repair. End-repaired samples were ligated to 3′ adapters, followed by enzymatic removal of excess adapter and reverse transcription. cDNA was PAGE purified (70 ~ 80 nt for ribosome profiling, 70 ~ 100 nt for mRNA-seq), circularized, and PCR-amplified. PCR products were further purified by AMPure (Beckman Coulter) and an 8% acrylamide gel. A Bioanalyzer (Agilent) was used for quantification to pool the samples. Pooled libraries were outsourced to Macrogen (Japan) and sequenced on a HiSeq 2500 (Illumina).

Sequenced read analysis

The adapter sequence was trimmed with cutadapt (*36*) and first mapped to the ribosomal DNA genome retrieved from Biomart martview (*37*). Fastq files after this step were deposited in Gene Expression Omnibus. Reads were then aligned to mouse (mm10) or human (hg19) transcripts using Bowtie 1.2.2 (*38*). These reference transcripts contain only CDSs of each gene and 30 nt upstream of CDSs, retrieved from Biomart martview. Multiple CDSs in a gene were removed and the longest CDS transcript was used. To validate ribosome profiling data, metagene analysis was performed. 5′ ends of ribosome footprints relative to translation start sites in all genes were accumulated and their sum at each position was normalized using averaged footprints.

Ribosome density calculation (validation of the noise reduction method)

To reduce noise, ribosome footprints at the same position of the same gene were converted to 1 count, no matter how many footprints were present at the same position (i.e., each position of each gene has 0 or 1 count, "binary count", of footprints). The robustness of this strategy was validated using publicly available ribosome profiling data of arginine-starved 293T cells (GSE113751) (*39*). In each gene, the relative distance of ribosomal P sites (11 nt from 5′ end of footprints) to indicated Arg codons was determined for each binary-converted ribosome footprint and ribosome densities of relative distance were accumulated. To normalize differences in total footprint numbers among samples, accumulated counts at each relative position were divided by averaged footprint counts between -50 to 200 nt relative to the corresponding Arg codons. These normalized counts with or without noise reduction were then plotted.

Ribosome density calculation (association of miRNA and ribosome density)

Focusing on miRNA expression greater than 15 FPKM in C2C12 mouse myoblast cells (GSE99399) (*40*), miRNA binding sites in mouse CDSs were predicted using the TarPmiR algorithm (*41*), using machine-learning approaches based on published CLASH data. As for miRNA binding sites in human CDSs, miRNAs greater than 10 RPM (GSE56836) (*42*) were selected. Publicly available CLASH data in HEK293 (*10*) (GSE46039) were used as well as publicly available ribosome profiling data in 293T cell with no pre-treatment with cycloheximide (GSE113751) (*39*), which confirm that the currently identified relation between ribosome footprints and miRNA binding sites is independent of cycloheximide pre-treatment. As indicated, miRNAs were categorized based on their binding strength. For random sequences, 10 random positions of each gene were selected and processed as described below.

We used the top 5,000 and 8,000 highly translated genes in mouse skeletal muscle and 293T cells, respectively. Ribosome footprints at the same position of the same gene were converted to 1 count. The relative distance between the 3′ end of ribosome footprints and the 5′ end of miRNA binding sites of each gene was determined for each binary-converted ribosome footprint. Then, ribosome densities of relative distance were accumulated. Accumulated counts at each position were divided by the sum of counts of all the positions to normalize differences in the number of total sequenced reads among samples. Normalized ribosome densities at each relative position were further normalized by similarly defined mRNA-seq densities.

Misfolding and aggregation assay

Cells were seeded 48 hr before harvest. Cells were treated with MG132 (Cayman Chemical) at a final concentration of 20 μM for the indicated time before harvest. For imaging, cells were fixed with 4% paraformaldehyde (Fujifilm, Japan) for 20 min, followed by a PBS wash and Hoechst 33342 staining (Dojindo, Japan). Images were acquired by confocal microscope TCS SP8 (Leica). For protein extraction, cells were washed with ice-cold PBS and lysed with lysis buffer (without DTT for IP) (20 mM Tris-HCl, 150 mM NaCl, 1 % Triton-X100, protease inhibitor cocktail without EDTA [Nacalai tesque, Japan], 10 % Glycerol, 13 unit/mL of Turbo DNase [Invitrogen], 1 mM DTT), followed by 30-min incubation on ice. Lysate was centrifuged at 1,000 g at 4 °C for 5 min. The supernatant, as total extract, was used for downstream experiments. To separate aggregated insoluble fraction, the supernatant was centrifuged at 20,000 g at 4 °C for 60 min. The pellet was washed with ice-cold PBS, followed by centrifugation at 20,000 g at 4 °C for 10 min. After discarding PBS, the aggregated insoluble fraction was dissolved in 2% SDS-containing buffer (20 mM Tris-HCl, 150 mM NaCl, 1% Triton-X100, protease inhibitor cocktail without EDTA, 10% glycerol, 1 mM DTT, 2 % SDS) for 20 min at room temperature.

A SUnSET assay (*43*) was used to quantify nascent protein synthesis rates. Cells were cultured in full medium with 5 μg/mL puromycin for 30 min. Cells were washed with ice-cold PBS and lysed with lysis buffer without DTT, followed by 30 min lysis on ice. To pellet the debris, lysate was centrifuged at 12,000 g at 4 °C for 5 min. Lysate was used to quantify total protein synthesis rates. To quantify nascent NBD1-reporter synthesis rates, immunoprecipitation against SBP (Streptavidin-Binding Peptide)-tag was performed. Lysate was incubated with Dynabeads M-280 Streptavidin (Invitrogen) overnight at 4 °C. Beads were washed with lysis buffer 5 times and dissolved in lysis buffer with DTT and 4x sample loading buffer (0.5 M Tris-HCl pH 6.8, 2 % SDS, 10% glycerol, bromophenol blue. DTT was added before use at a final concentration of 25 mM).

Total extract and insoluble proteins were quantified with a Pierce BCA protein assay kit (Thermo Scientific) so as to load equal amounts and incubated for 10 min at 70 °C for western blot. NBD1 and mCherry were detected with anti-SBP and anti-Flag antibodies, respectively. For quantification, insoluble fractions of NBD1 were normalized against γTUBULIN instead of mCherry to avoid biased quantification. This is because the relative amount of insoluble mCherry in sgDROSHA decreased compared to NTC (Fig. S2G), which positively biases the results. Image Studio Lite (LI-COR Biosciences) was used for quantification. Antibodies used were as follows: anti-Flag (F7425, Sigma), anti-SBP (sc-101595, Santa Cruz), anti-AGO1 (07-599, Millipore), anti-AGO2 (2897S, Cell Signaling), anti-DROSHA (sc-393591, Santa Cruz), anti-Puromycin (EQ0001, Kerafast), anti-HSC70 (sc-24, Santa Cruz), anti-γTUBULIN (T3320, Sigma), anti-GAPDH (2118S, Cell Signaling), anti-Rabbit IgG (NA9340, GE Healthcare), mouse-IgGk BP-HRP (sc-516102, Santa Cruz). As for miRNA-deficient cells and shRNA-expressing cells, experiments from infection to insoluble NBD1 assays were repeated at least 5 times with similar results. Other experiments were repeated at least 3 times with similar results.

RIP-qPCR

Cells were crosslinked with medium with 2.5 mM DSP (Dojindo) for 45 seconds, followed by incubation of medium with 50 mM Tris-HCl to quench the reaction. Crosslinked cells were washed with ice-cold PBS with 100 μg/mL CHX and lysed with lysis buffer (20 mM Tris-HCl, 150 mM NaCl, 1% Triton-X100, 0.1% SDS, protease inhibitor cocktail without EDTA, 10% glycerol, 13 unit/mL of Turbo DNase, 5 mM $MgCl_2$, 100 μg/mL CHX, and 1% SuperaseIn). Anti-HSC70 (sc-24, Santa Cruz) was incubated with pre-washed Dynabeads M-280 sheep anti-mouse IgG (Invitrogen) in lysis buffer for 4 hr at 4 °C, followed by a brief wash and incubation with cell lysate overnight at 4 °C. Then, beads were washed with high-salt buffer (20 mM Tris-HCl, 600 mM NaCl, 1% Triton-X100, 0.1% SDS, 5 mM $MgCl_2$, 100 μg/mL CHX, and 1% SuperaseIn) three times, followed by three washes with lysis buffer. Protein was eluted with lysis buffer and 4x sample buffer with DTT at 70 °C for 10 min. RNA was extracted by adding ISOGEN II (Nippongene, Japan) to beads, followed by standard isopropanol-based precipitation with GlycoBlue (Invitrogen).

All extracted RNA was reverse transcribed using PrimeScript II (Takara Bio, Japan) and real time qPCR was carried out with TB Green Premix Ex Taq II (Takara Bio) and ViiA 7 (Thermo Fisher Scientific). *NBD1 reporter* expression was quantified by targeting the SBP tag sequence and normalized against *γTUBULIN* or *GAPDH* expression.

30. Y. Kato, S. Miyaki, S. Yokoyama, S. Omori, A. Inoue, M. Horiuchi, H. Asahara, Real-time functional imaging for monitoring miR-133 during myogenic differentiation. *Int. J. Biochem. Cell Biol.* **41**, 2225–2231 (2009).
31. N. E. Sanjana, O. Shalem, F. Zhang, Improved vectors and genome-wide libraries for CRISPR screening. *Nat. Methods*. **11**, 783–784 (2014).
32. S. A. Stewart, Lentivirus-delivered stable gene silencing by RNAi in primary cells. *RNA*. **9**, 493–501 (2003).
33. R. J. Golden, B. Chen, T. Li, J. Braun, H. Manjunath, X. Chen, J. Wu, V. Schmid, T.-C. Chang, F. Kopp, A. Ramirez-Martinez, V. S. Tagliabracci, Z. J. Chen, Y. Xie, J. T. Mendell, An Argonaute phosphorylation cycle promotes microRNA-mediated silencing. *Nature*. **542**, 197–202 (2017).
34. N. T. Ingolia, S. Ghaemmaghami, J. R. S. Newman, J. S. Weissman, Genome-Wide Analysis in Vivo of Translation with Nucleotide Resolution Using Ribosome Profiling. *Science*. **324**, 218–223 (2009).
35. H. Sako, K. Yada, K. Suzuki, Genome-Wide Analysis of Acute Endurance Exercise-Induced Translational Regulation in Mouse Skeletal Muscle. *PLOS ONE*. **11**, e0148311 (2016).
36. M. Martin, Cutadapt Removes Adapter Sequences From High-Throughput Sequencing Reads, *EMBnet J.* **17**, 10-12 (2011).
37. D. Smedley, S. Haider, S. Durinck, L. Pandini, P. Provero, J. Allen, O. Arnaiz, M. H. Awedh, R. Baldock, G. Barbiera, P. Bardou, T. Beck, A. Blake, M. Bonierbale, A. J. Brookes, G. Bucci, I. Buetti, S. Burge, C. Cabau, J. W. Carlson, C. Chelala, C. Chrysostomou, D. Cittaro, O. Collin, R. Cordova, R. J. Cutts, E. Dassi, A. D. Genova, A. Djari, A. Esposito, H. Estrella, E. Eyras, J. Fernandez-Banet, S. Forbes, R. C. Free, T. Fujisawa, E. Gadaleta, J. M. Garcia-Manteiga, D. Goodstein, K. Gray, J. A. Guerra-

Assunção, B. Haggarty, D.-J. Han, B. W. Han, T. Harris, J. Harshbarger, R. K. Hastings, R. D. Hayes, C. Hoede, S. Hu, Z.-L. Hu, L. Hutchins, Z. Kan, H. Kawaji, A. Keliet, A. Kerhornou, S. Kim, R. Kinsella, C. Klopp, L. Kong, D. Lawson, D. Lazarevic, J.-H. Lee, T. Letellier, C.-Y. Li, P. Lio, C.-J. Liu, J. Luo, A. Maass, J. Mariette, T. Maurel, S. Merella, A. M. Mohamed, F. Moreews, I. Nabihoudine, N. Ndegwa, C. Noirot, C. Perez-Llamas, M. Primig, A. Quattrone, H. Quesneville, D. Rambaldi, J. Reecy, M. Riba, S. Rosanoff, A. A. Saddiq, E. Salas, O. Sallou, R. Shepherd, R. Simon, L. Sperling, W. Spooner, D. M. Staines, D. Steinbach, K. Stone, E. Stupka, J. W. Teague, A. Z. Dayem Ullah, J. Wang, D. Ware, M. Wong-Erasmus, K. Youens-Clark, A. Zadissa, S.-J. Zhang, A. Kasprzyk, The BioMart community portal: an innovative alternative to large, centralized data repositories. *Nucleic Acids Res.* **43**, W589–W598 (2015).
38. B. Langmead, C. Trapnell, M. Pop, S. L. Salzberg, Ultrafast and memory-efficient alignment of short DNA sequences to the human genome. *Genome Biol.* **10**, R25 (2009).
39. A. M. Darnell, A. R. Subramaniam, E. K. O'Shea, Translational Control through Differential Ribosome Pausing during Amino Acid Limitation in Mammalian Cells. *Mol. Cell*. **71**, 229-243.e11 (2018).
40. K. He, G. Wu, W.-X. Li, D. Guan, W. Lv, M. Gong, S. Ye, A. Lu, A transcriptomic study of myogenic differentiation under the overexpression of PPARγ by RNA-Seq. *Sci. Rep.* **7**, 15308 (2017).
41. J. Ding, X. Li, H. Hu, TarPmiR: a new approach for microRNA target site prediction. *Bioinformatics*. **32**, 2768–2775 (2016).
42. H. P. Bogerd, A. W. Whisnant, E. M. Kennedy, O. Flores, B. R. Cullen, Derivation and characterization of Dicer- and microRNA-deficient human cells. *RNA*. **20**, 923–937 (2014).
43. E. K. Schmidt, G. Clavarino, M. Ceppi, P. Pierre, SUnSET, a nonradioactive method to monitor protein synthesis. *Nat. Methods*. **6**, 275–277 (2009).

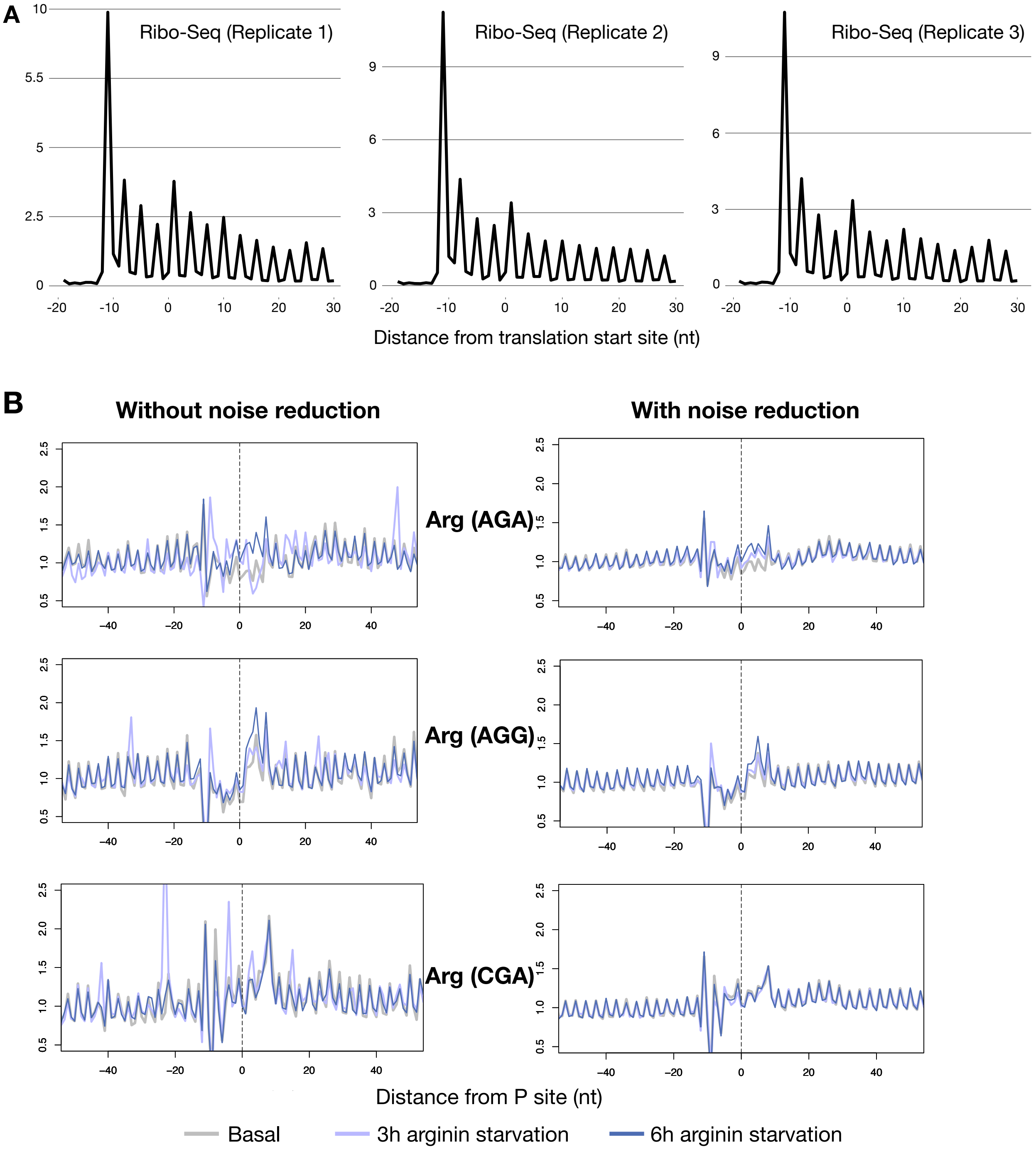

A
Ribo-Seq (Replicate 1)
Ribo-Seq (Replicate 2)
Ribo-Seq (Replicate 3)
Distance from translation start site (nt)
B
Without noise reduction
With noise reduction
Arg (AGA)
Arg (AGG)
Arg (CGA)
Distance from P site (nt)
Basal
3h arginin starvation
6h arginin starvation

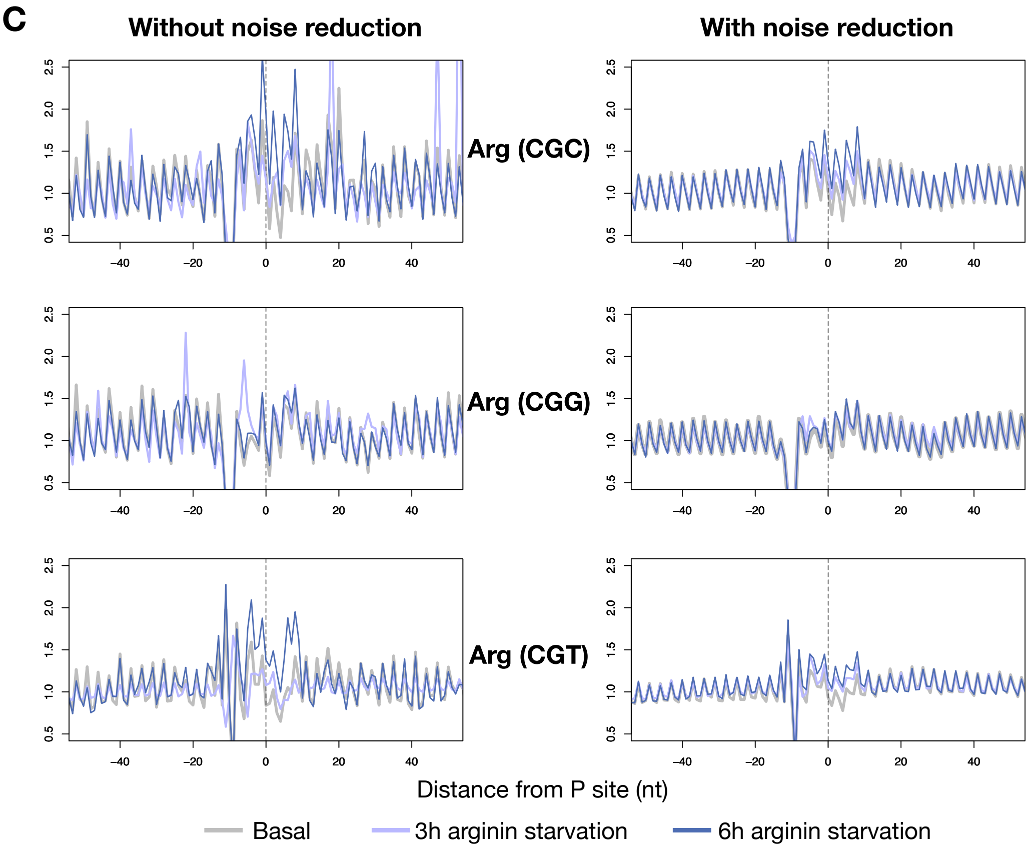


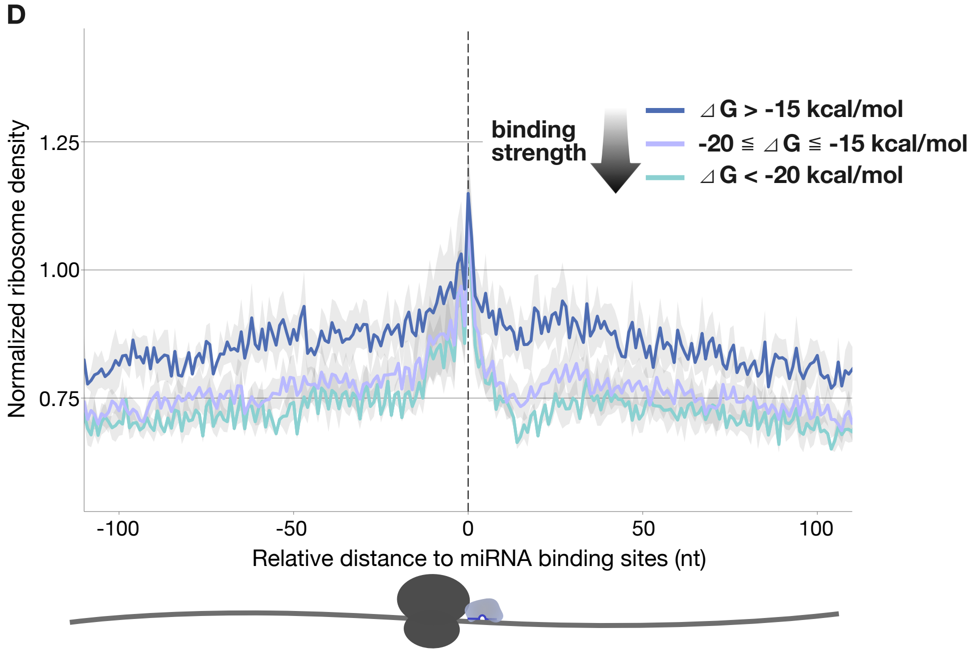


**Fig. S1.** (A) Meta-gene analysis of ribosome profiling in mouse skeletal muscle. (B, C) Validation of the noise-reduction method used in this study (Fig. 1B, S1D). Before and after noise reduction, densities of ribosomal P sites relative to the indicated Arg codons were plotted under basal, 3 hr, or 6 hr of Arg depletion. Larger ribosomal peaks at ~ +3 nt (ribosomal A site) are seen in a time-dependent manner compared to the basal condition. (D) Ribosome densities normalized by total sequenced reads are shown using publicly available 293T ribosome profiling data and CLASH data.

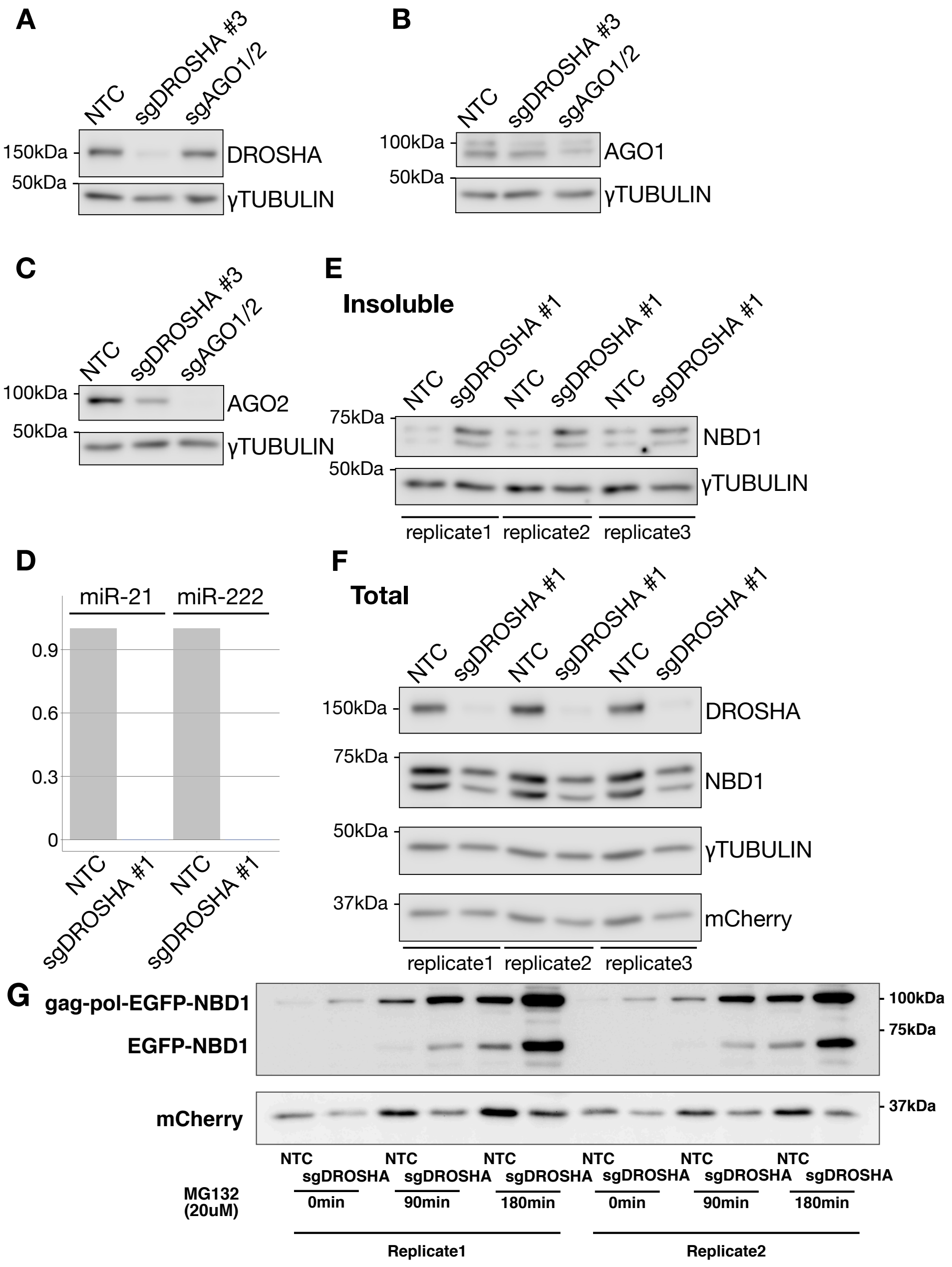


**Fig. S2.** (A, B, C) Knockout efficiency of each knockout cell used in Figs. 2C and D. All are non-clonal cells. (D) Quantification of miRNA expression with TaqMan assays, normalized against U6. (E) Immunoblots of insoluble protein fractions of DROSHA knockout with a different guide sequence after 120 min of MG132 (20 μM) treatment. (F) Total extract of (E). As in (E) the NBD1 blot shows two bands, cleaved by 1st P2A or 2nd P2A sequences. (G) Time-dependent accumulation of gag-pol-NBD1 reporter aggregates in insoluble fractions. The reporter used here expresses both gag-pol-fused NBD1 and gag-pol-free NBD1 as indicated gag-pol-EGFP-NBD1 and EGFP-NBD1, respectively. The guide sequence used to knockout DROSHA is the same as in Figs. S2E and F. The relative amount of mCherry in sgDROSHA decreased compared to NTC, which leads to positive bias if NBD1 is normalized against mCherry. Therefore, insoluble NBD1 was normalized against γTUBULIN.

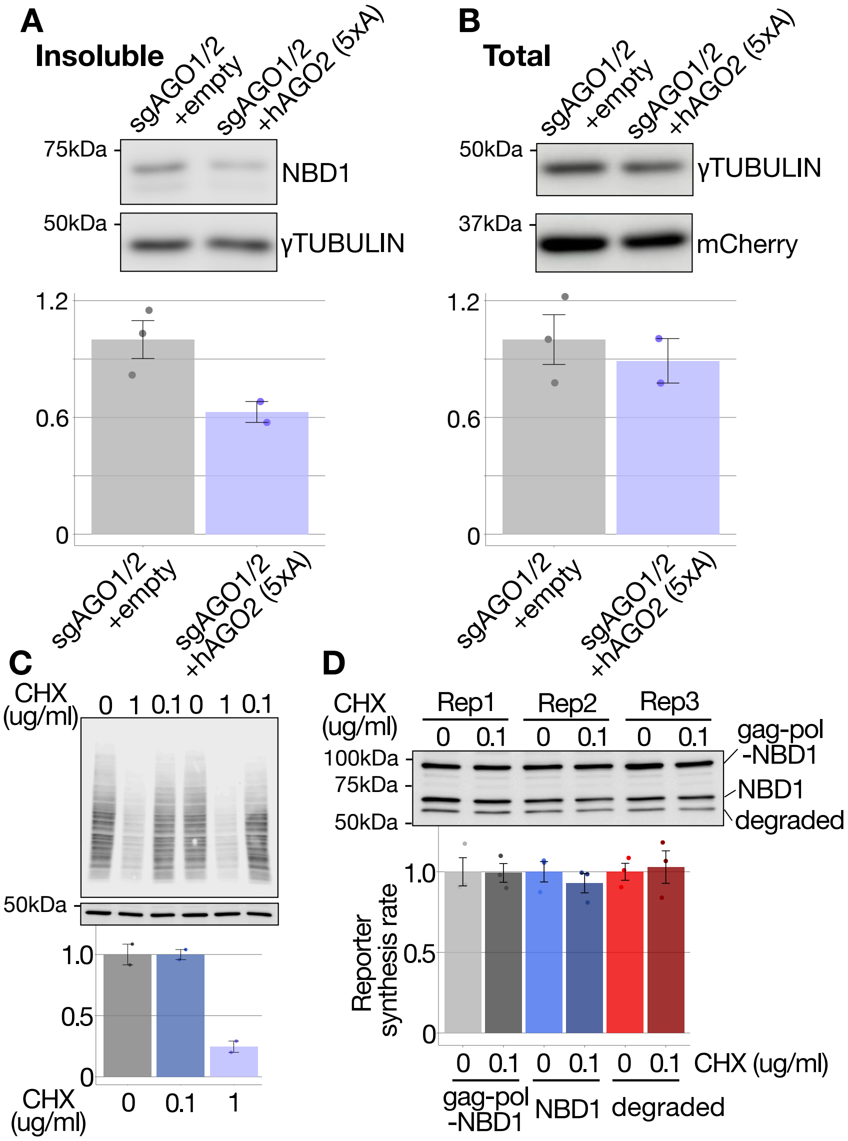


**Fig. S3.** (A, B) AGO2 (5xA) over-expression rescues sgAGO1/2-induced aggregation. Immunoblots and quantification of insoluble (A) and total (B) protein fractions. (C, D) SUnSET assay for total (C) and reporter-specific (D) protein synthesis. A gag-pol-NBD1 reporter used here expresses both gag-pol-fused NBD1 and gag-pol-free NBD1 as in Fig. S2G. Total protein synthesis rates detected with anti-puromycin were normalized against γTUBULIN for the plot.

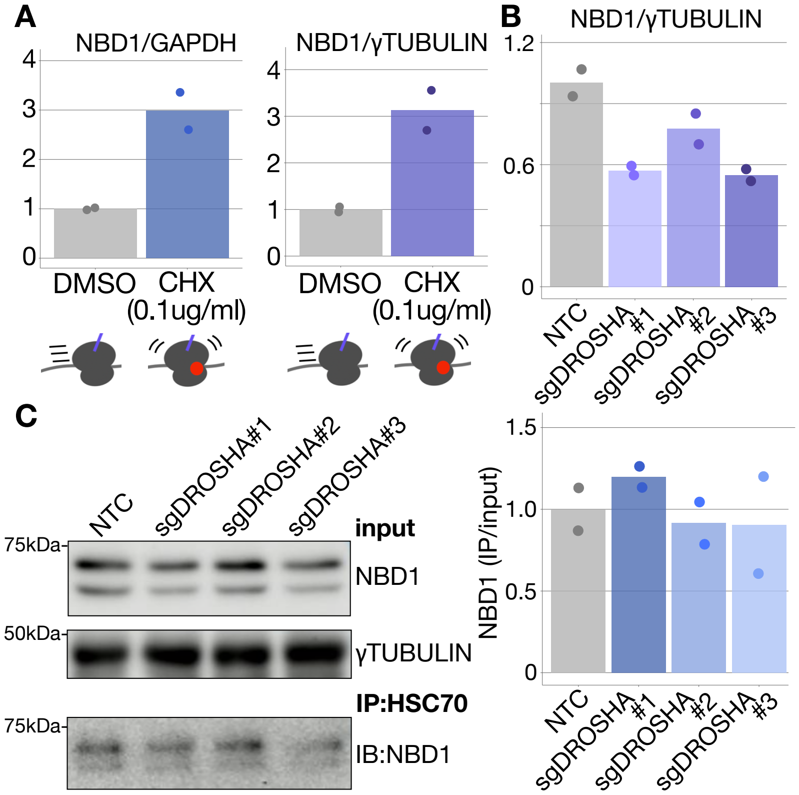


**Fig. S4.** (A) Validation of the RIP-qPCR method in this study. Cells were first treated with CHX (0.1 μg/mL) for 60 min, followed by crosslinking. CHX (0.1 μg/mL) slows ribosomal elongation, which co-translationally recruit more HSC70 compared to DMSO only. *NBD1 reporter* mRNA levels were normalized against *GAPDH* or *γTUBULIN*. IP: HSC70. (B) RIP-qPCR as in Fig 4B. Quantification of *NBD1 reporter* mRNA levels, normalized against *γTUBULIN*. (C) HSC70 recruitment efficiency for mature NBD1 reporter was determined. NBD1 reporter abundance detected in IP (HSC70) was normalized by the reporter abundance in γTUBULIN-normalized input and quantified.

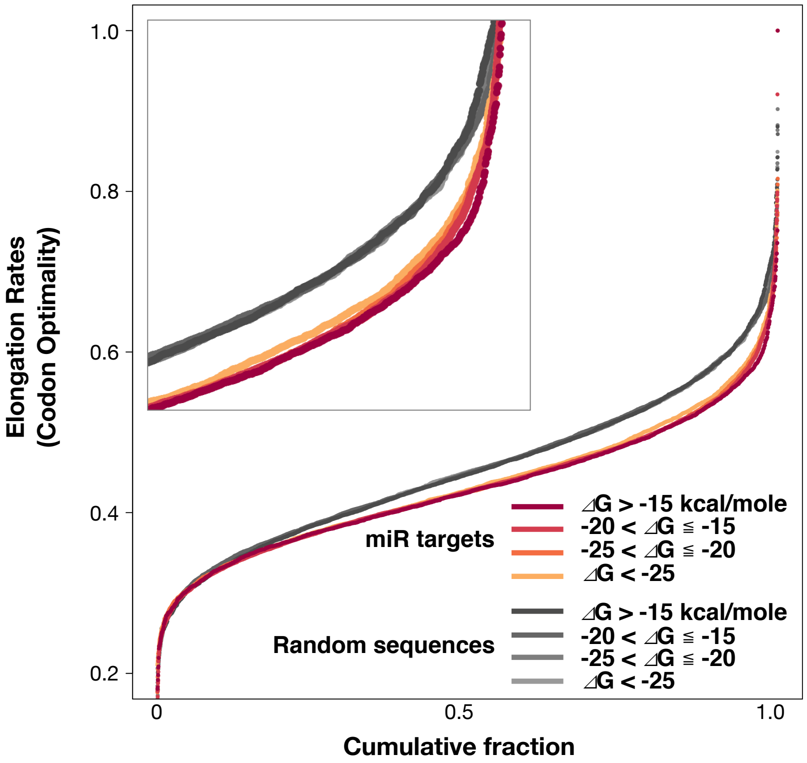


**Fig. S5.** Association of miRNA binding strength and corresponding codon optimality. Codon optimality scores were obtained from (*3*). miRNA target sequences are more likely to have lower codon optimality, making it more difficult to design NBD1 mutations without affecting codon optimality, RNA secondary structure, or RNA folding energy to alter miRNA binding.

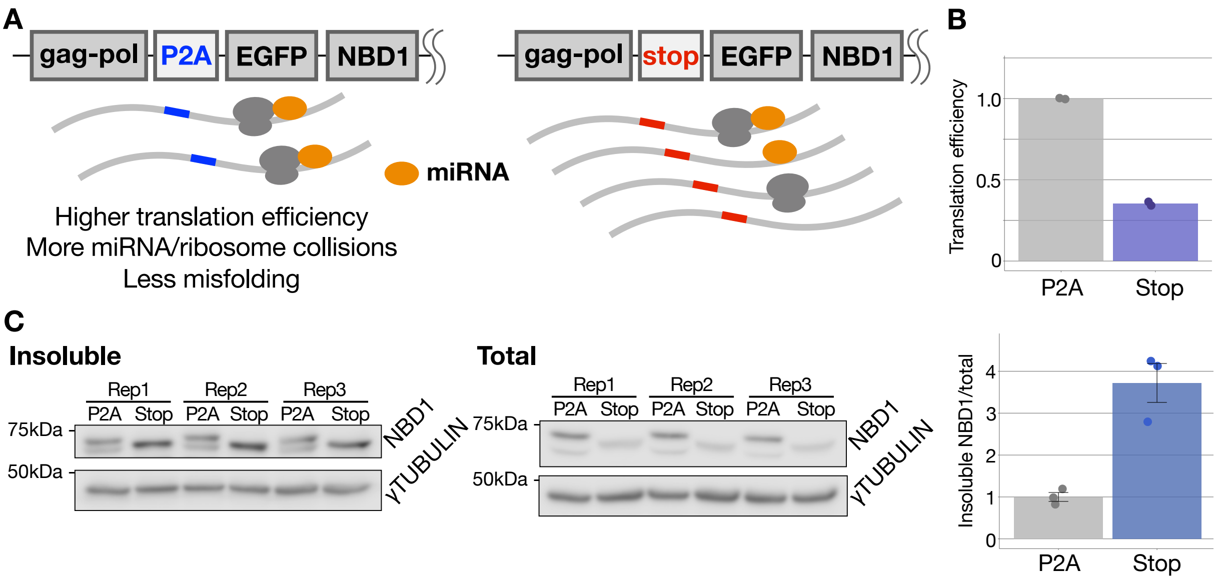


**Fig. S6.** (A) Schematic image of two reporters (P2A and stop-NBD1) to cause different translation efficiencies. In the P2A reporter (equal to NBD1-reporter), ribosomes that initiate translation of gag-pol continue translating EGFP-NBD1; however, in the stop-NBD1 reporter, only ribosomes that fail to recognize the gag-pol translation initiation site can initiate EGFP-NBD1 translation. This causes low translation efficiency in the stop-NBD1 reporter compared to P2A (NBD1-reporter). Since ribosome-miRNA collision rates are assumed to be low if translation efficiency is low, given our working model, more misfolding relative to total reporter abundance is expected. (B) Translation efficiency of the P2A (NBD1) and stop-NBD1 reporter. Efficiency was calculated by ribosome density (ribosome profiling) divided by mRNA abundance (mRNA-seq). The difference was mainly caused by the difference in mRNA abundance (P2A : Stop = 1 : 3.60) rather than ribosome density (P2A : Stop = 1 : 1.27). (C) Immunoblots and quantification of relative reporter abundance. P2A (NBD1-reporter) and stop-NBD1 reporter were normalized against γTUBULIN in both insoluble and total fractions. Then, insoluble fractions of each reporter were normalized by that in total fractions for this plot.

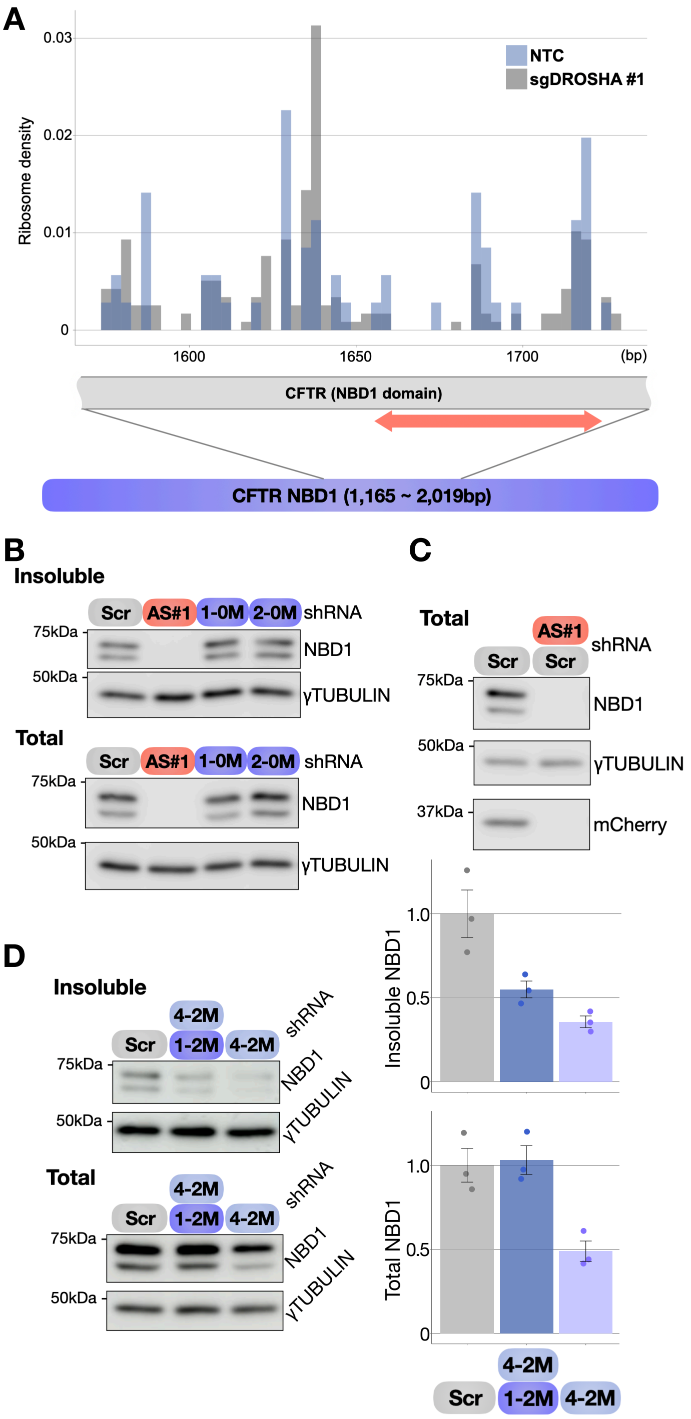


**Fig. S7.** (A) Ribosome densities of the NBD1 domain in NTC and sgDROSHA are shown. Replicate of Fig. 5A. Raw ribosomal footprint counts were normalized against total footprint numbers aligned to NBD1 domain sequence. Bin = 3 nt, showing single-codon resolution. The target (~ 60 nt) of shRNA is indicated by a red arrow. (B) Immunoblots of insoluble fractions of NBD1 and total NBD1 after 150 min of MG132 (20 μM) treatment. Fully complementary antisense shRNA (AS#1) and non-cleaving shRNA. shRNA "1-0M" has 2 mismatches at the $10^{th}$ and $11^{th}$ nucleotides of AS#1. Scr is scramble control. 2-0M targets outside of the 60-nt region. (C) Immunoblot validating functionality of tandemly expressed shRNA. (D) Immunoblots and quantification of insoluble fractions of NBD1 and total NBD1 after 150 min of MG132 (20 μM) treatment.

Table S1. Names and sequences used in this study.

| Names | Sequence (5´ to 3´) |
|---|---|
| CFTR_NBD1_Fw | ATATActcgagGGAGGCGGAGGCTCTACTACAGAAGTAGTGATGGAGAATGTAA |
| CFTR_NBD1_Rv | ATccgcggTCCTTCTAATGAGAAACGGTGTAAGGT |
| SBP_Fw | ATGGATGAGAAGACAACTGGGT |
| SBP_Rv | GCTCTCTTTGACCTTGGGGAT |
| human_GAPDH_Fw | GCACCGTCAAGGCTGAGAAC |
| human_GAPDH_Rv | ATGGTGGTGAAGACGCCAGT |
| human_gTubulin_Fw | AGGAAGTCTCCCTACCTGCC |
| human_gTubulin_Rv | AGGTTCTCTCGAAGAGCGAGG |
| NTC_oligo1 | caccgGTATTACTGATATTGGTGGG |
| NTC_oligo2 | aaacCCCACCAATATCAGTAATACc |
| sgDROSHA_#1_oligo1 | caccgTTGGTGGTAGCGGATATGAT |
| sgDROSHA_#1_oligo2 | aaacATCATATCCGCTACCACCAAc |
| sgDROSHA_#2_oligo1 | caccgAATGAGACGAGAAGTAACGG |
| sgDROSHA_#2_oligo2 | aaacCCGTTACTTCTCGTCTCATTc |
| sgDROSHA_#3_oligo1 | caccgGCATTAGGCATTGGTGGTAG |
| sgDROSHA_#3_oligo2 | aaacCTACCACCAATGCCTAATGCc |
| sgAGO1_oligo1 | caccgACAGACTGTGGAGTGCACAG |
| sgAGO1_oligo2 | aaacCTGTGCACTCCACAGTCTGTc |
| sgAGO2_oligo1 | caccgGCAGACGGTGGAGTGCA |
| sgAGO2_oligo2 | aaacTGCACTCCACCGTCTGCc |
| Scr | CCTAAGGTTAAGTCGCCCTCGcgaaCGAGGGCGACTTAACCTTAGG |
| AS#1 | GCAAGAATTTCTTTAGCAAGAcgaaTCTTGCTAAAGAAATTCTTGC |
| 1-0M | GCAAGAATTTgaTTAGCAAGAcgaaTCTTGCTAAtcAAATTCTTGC |
| 1-1M | cCAAGAATTTgaTTAGCAAGAcgaaTCTTGCTAAtcAAATTCTTGg |
| 1-2M | cgAAGAATTTgaTTAGCAAGAcgaaTCTTGCTAAtcAAATTCTTcg |
| 2-0M | GGGATGTGATagTTTCGACCAcgaaTGGTCGAAActATCACATCCC |
| 3-0M | TACCTAGATGTaaTAACAGAAAcgaaTTTCTGTTAttACATCTAGGTA |
| 4-0M | ATACAAAGATcgTGATTTGTAcgaaTACAAATCAcgATCTTTGTAT |
| 4-2M | cgACAAAGATcgTGATTTGTAcgaaTACAAATCAcgATCTTTGTcg |
